# Epigenome-wide association study of placental DNA methylation and maternal exposure to night shift work in the Rhode Island Child Health Study

**DOI:** 10.1101/265777

**Authors:** Danielle A. Clarkson-Townsend, Todd M. Everson, Maya A. Deyssenroth, Amber A. Burt, Karen E. Hermetz, Ke Hao, Jia Chen, Carmen J. Marsit

**Affiliations:** Department of Environmental Health, Rollins School of Public Health, Emory University, Atlanta, GA, USA; Department of Environmental Medicine and Public Health, Icahn School of Medicine at Mount Sinai, New York, NY, USA; Department of Genetics and Genomic Sciences, Icahn School of Medicine at Mount Sinai, New York, NY, USA

**Keywords:** Night shift work, epigenetics, epigenome-wide association study, neuron navigator 1, circadian disruption

## Abstract

**Objectives:** Circadian disruption from environmental and occupational exposures can potentially impact health, including offspring health, through epigenetic alterations. Night shift workers experience circadian disruption, but little is known about how this exposure could influence the epigenome of the placenta, which is situated at the maternal-fetal interface. To investigate whether night shift work is associated with variations in DNA methylation patterns of placental tissue, we conducted an epigenome-wide association study (EWAS) of night shift work.

**Methods:** CpG specific methylation genome-wide of placental tissue (measured with the Illumina 450K array) from participants (n=237) in the Rhode Island Child Health Study (RICHS) who did (n=53) and did not (n=184) report working the night shift was compared using robust linear modeling, adjusting for maternal age, pre-pregnancy smoking, infant sex, maternal adversity, and putative cell mixture.

**Results:** Night shift work was associated with differential methylation in placental tissue, including CpG sites in the genes *NAV1, SMPD1, TAPBP, CLEC16A, DIP2C, FAM172A*, and *PLEKHG6* (Bonferroni-adjusted p<0.05). CpG sites within *NAV1, MXRA8, GABRG1, PRDM16, WNT5A*, and *FOXG1* exhibited the most hypomethylation, while CpG sites within *TDO2, ADAMTSL3, DLX2*, and *SERPINA1* exhibited the most hypermethylation (BH q<0.10). PER1 was the only core circadian gene demonstrating differential methylation. Functional analysis indicated GO-terms associated with cell-cell adhesion.

**Conclusions:** Night shift work was associated with differential methylation of the placenta, which may have implications for fetal health and development. Additionally, neuron navigator 1 (NAV1) may play a role in the development of the human circadian system.

**What is already known about this subject?:** Night shift work and circadian disruption may play a role in the development and progression of many diseases. However, little is known about how circadian disruption impacts human fetal health and development.

**What are the new findings?:** Working the night shift is associated with altered placental methylation patterns, and particularly, neuron navigator 1 (NAV1) may play a role in the development of the human circadian system.

**How might this impact on policy or clinical practice in the foreseeable future?:** Night shift work prior to or during pregnancy may alter the placental epigenome, which has implications for fetal health. Further studies are needed to evaluate night shift work as a possible risk factor for gestational diabetes and to evaluate the impact of circadian disruption on fetal health and development.

## INTRODUCTION

Disruption of circadian rhythms is an occupational hazard for people who work the night shift and is associated with negative health outcomes such as cancer, metabolic disorders, and neurological disorders ^1^. Animal models also demonstrate altered metabolism, hormonal signaling, body temperature rhythms, and adiposity in experiments mimicking shift work exposure ^2^. The health risks posed by night shift work may have large public health consequences, as approximately 15% of American employees work outside of the traditional 9AM-5PM work schedule ^3^. While some aspects of the circadian system may return to normal after a regular schedule of night shift work, studies suggest the majority of regular night shift workers (~97%) aren’t able to fully adapt their endogenous circadian rhythms to their work schedules ^4^. People also commonly experience circadian disruption when exposed to light at night (LAN) ^5^.

The daily and seasonal patterns of light and dark exposure are an important environmental stimulus. Organisms have evolved an internal timekeeping mechanism, the circadian clock, to generate rhythms of biological activity to adapt to predictable environmental changes, increasing physiological efficiency and fitness. The core circadian clock consists of feedback loops of transcription factors (TF) that generate oscillating cycles of gene transcription and translation. These endogenously generated rhythms rely on cues, such as light, to synchronize patterns of physiological activity with the external environment. When light enters the eye, it activates visual photopigment in photoreceptors and in melanopsin-containing retinal ganglion cells (mRGCs). The light signal is transmitted via the retinohypothalamic tract to the suprachiasmatic nucleus (SCN) of the hypothalamus, the "master clock” that sets the body’s peripheral clocks ^6^. This elegant system of interdependent signaling ensures processes such as protein synthesis, fatty acid metabolism, and insulin release occur at the appropriate times to adapt to daily and seasonal changes in the environment ^7^. However, when circadian rhythms are misaligned, cell signaling becomes inefficient and dissonant, contributing to disease.

Animal studies suggest circadian disruption *in utero* negatively affects the health and development of offspring ^8^. Mice exposed to a 22-hour light-dark cycle, instead of the normal 24-hour cycle, had altered methylation patterns in the SCN and altered circadian behavior; differential methylation was also found for genes related to axonal migration, synaptogenesis, and neuroendocrine hormones ^9^. Additionally, chronic changes in the photoperiod of pregnant rats caused increased leptin levels, insulin secretion, fat deposition, and decreased glucose tolerance of offspring in adulthood ^10^. Little is known about the impact of light or circadian rhythms in human pregnancy or on long-term fetal programming, although there appear to be only small risks of negative reproductive health outcomes associated with shift work ^11^. During pregnancy, the placenta acts as a mediator between the maternal and fetal environment to regulate growth and development; yet, little attention has been paid to the impact of circadian disruption on placental function. Variation in DNA methylation patterns in the placenta can affect placental function. Because the placenta is composed of fetal DNA, methylation of placental tissue may reflect fetal exposures and future health effects. In this study, we conducted an epigenome-wide association study (EWAS) to investigate whether night shift work is associated with differences in DNA methylation in the placental epigenome, which can impact long-term health outcomes in the offspring.

## METHODS

### Study population - The Rhode Island Child Health Study (RICHS)

RICHS is a hospital-based cohort study of mothers and infants in Rhode Island, described in detail in ^12^. Briefly, from 2009 to 2014, 844 women between the ages of 18-40 and their infants were enrolled at the Women and Infants Hospital of Rhode Island, oversampling for large and small for gestational age (LGA, SGA, respectively) infants and matching each to an appropriate for gestational age (AGA) control by maternal age ( ±2 years), sex, and gestational age ( ± 3 days). RICHS enrolled only full-term (≥37 weeks), singleton deliveries without congenital or chromosomal abnormalities. Demographic information was collected from a questionnaire administered by a trained interviewer and clinical outcome information was obtained from medical records. Information on night shift work was obtained from questionnaire by first asking, "Have you ever worked outside the home? (Yes/No)" and if "Yes”, participants were asked "If yes, please list all of the jobs you have had starting with your current job first. Please indicate whether you worked a swing shift or a night shift on any of these jobs”. To indicate shift jobs, the questionnaire included check boxes for "Yes” and "No” under a category for "Night Shift”. For this analysis, only the most recently reported job history was used. To adjust for socioeconomic factors while avoiding multicollinearity, we used an adversity score index to adjust for household income, maternal education, marital status and partner support. The cumulative risk score ranged from 0 to 4, with 0 representing the lowest level of adversity and 4 representing the highest level of adversity. A higher risk score was given to women whose median household income (adjusting for the number of people in the household) fell below the federal poverty line for the year the infant was born (+1), to women whose household was larger than 6 (+1), to women who were single and did not receive support from a partner (+1) and to women whose highest level of education was high school or less (+1) ^13^.

### Placental sample collection and measurement of DNA methylation

Genome-wide DNA methylation arrays were obtained on 334 placentae parenchyma samples in RICHS as previously described ^14^. The QA/QC process has been described elsewhere ^15^, including functional normalization, BMIQ, and ′ComBAT′ to adjust for technical variations and batch effects in R ^14 16^. Briefly, we used the ′minfi′ package in R to convert the raw methylation files to β values, a ratio of methylation ranging from 0 to 1, for analysis. Probes associated with the X or Y chromosomes, SNP-associated (within 10bp of the target CpG and with minor allele frequency >1%), identified as cross-reactive or polymorphic by Chen et al ^17^, or with poor detection p-values were excluded, yielding 334,692 probes for analysis in this study ^14^. DNA methylation array data for RICHS can be found in the NCBI Gene Expression Omnibus (GEO) database with the number GSE75248. Women with missing information on pre-pregnancy smoking status (“No”/“Yes”), defined as smoking 3 months prior to pregnancy, or adversity score were not included in the analysis. Women who did not provide an answer for the nightshift variable (n=16) were recoded to “No”. This study included the 237 mother-infant pairs within RICHS for which gene methylation data and the necessary demographic information were available.

### Placental RNA sequencing

Gene expression was measured using the Illumina HiSeq 2500 system in 199 placental samples from RICHS; methods have been previously described ^18^. After standard QA/QC procedures, final data were normalized to log2 counts per million (logCPM) values. Raw data is available in the NCBI sequence read archive (SRP095910).

### Statistical analyses

Because a reference panel for placental cell types does not yet exist, we used a validated reference-free method, the ′RefFreeEWAS′ package in R, to adjust for heterogeneity in cell-type composition ^19 20^. We implemented the RefFree estimation via the same process described in detail in our lab’s prior work ^21^, and identified 8 components to represent the putative cell mixture in our placental samples. We also examined the outlier screening plots of the cell mixture array for extreme outliers. We then conducted an EWAS using robust linear modeling by regressing CpG methylation β-values on night shift work (“No”/”Yes”), adjusting for putative cell mixture, maternal age (years), pre-pregnancy smoking status (“No”/”Yes”) adversity score (0-4)^22^, and sex of the infant (“Female”/”Male”). To adjust for multiple comparisons, we used the Bonferroni method and the Benjamini and Hochberg (BH) false discovery rate (FDR) methods. To evaluate the extent of *in utero* night shift exposure, we compared job and delivery date data. A secondary analysis using data from women who provided night shift job information (n=221) without recoding missing to “No” was also performed. While the distributions of gestational diabetes mellitus (GDM) and BMI differed between non-night shift and night shift workers, they were not adjusted for because they may be part of the causal pathway; numerous studies have found night shift work is associated with the development of obesity and metabolic diseases ^23 24^. We also investigated differentially methylated regions (DMRs) using the ′Bumphunter′ package in R ^25^. We modeled the β-values between non-night shift workers and night shift workers, controlling for the same variables as the EWAS. CpG sites within 500 base pairs were clustered together and β-values were modeled against a null distribution generated via bootstrapping; sites with differential methylation of 2% or more were considered to be possible DMRs.

To examine the functional implications of night shift work-associated DNA methylation (BH q<0.05), we also conducted an expression quantitative trait (eQTM) analysis using ′MEAL′ in R to investigate whether methylation was associated with gene expression in the RICHS samples with methylation and expression data available (n=199). Using robust linear modeling, we regressed the expression levels of genes within a 100kb window of the CpG site on methylation β-values (p<0.05).

### Bioinformatic analyses

To better understand the biological significance of the EWAS results, we analyzed the association of the top 298 CpG sites (BH q<0.10) with GO-terms and KEGG pathway enrichment in R using the ′missMethyl′ package ^26^. We analyzed the top 298 sites (BH q<0.10) because enrichment analyses generally require a few hundred genes to determine common pathways. We also evaluated genes of CpG sites with BH q<0.05 for rhythmicity with the CircaDB database ^27^. We analyzed all CIRCA experimental data using the JTK filter with a q- value probability cut-off of 0.05 and a JTK phase range of 0-40 ^28^. To investigate whether the EWAS results were associated with previous GWAS findings, the genes of CpG sites and DMRs that were significant after Bonferroni adjustment (p<0.05) were compared to gene results (p<5×10^−8^) in the GWAS catalog of the National Human Genome Research Institute and the European Bioinformatics Institute (NHGRI-EBI) ^29^.

## RESULTS

### Demographic and medical information

Demographic information for the women included in the methylation analysis is provided in Table 1. Comparing results from women who provided night shift job information (n=221) without recoding missing to “No” did not indicate any large differences in demographic features (Tables S1). Overall, women who reported working the night shift were more likely to be younger, smokers pre-pregnancy, cases of GDM, single and never married, lower household income, and higher adversity (p<0.05). While not statistically significant, women who worked the night shift trended towards a higher BMI and an evening chronotype. Of those included in the analysis, one participant reported taking melatonin and she was not a night shift worker. Additionally, 37 out of the 53 (70%) night shift workers reported working the night shift during pregnancy.

**Table 1.**
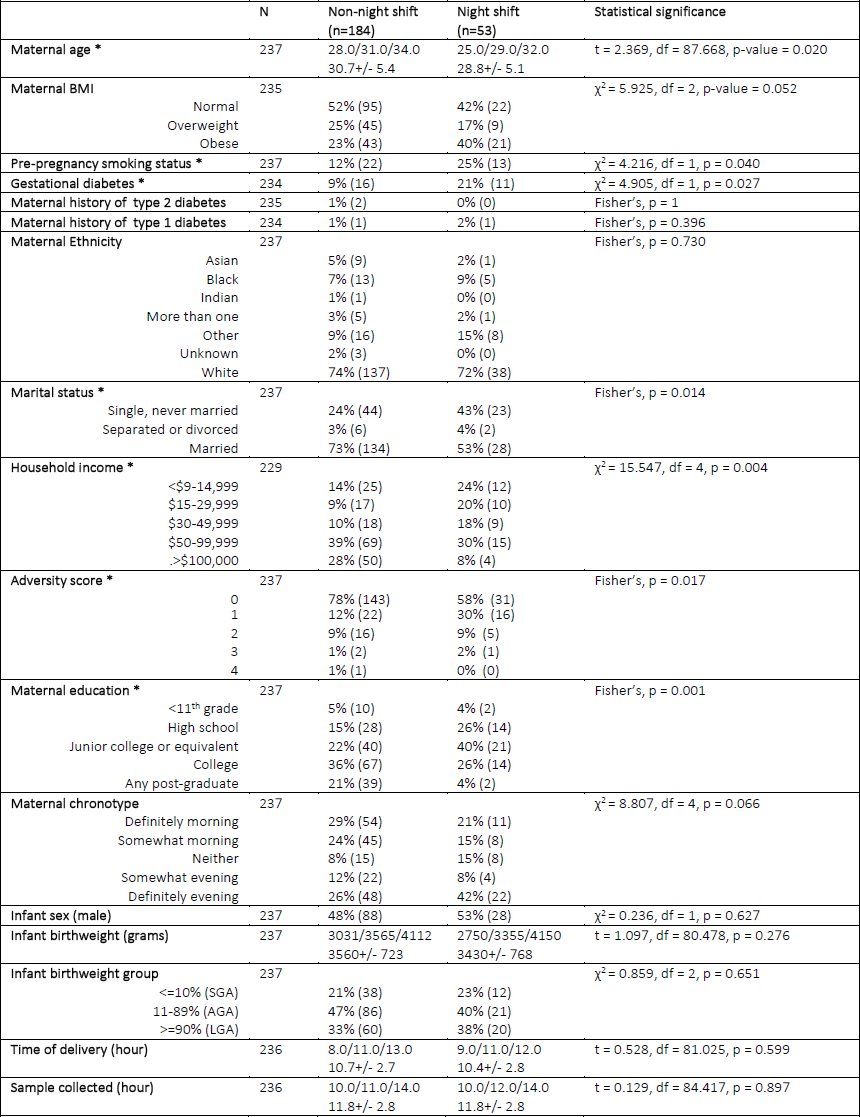
Demographic characteristics of those included in the EWAS methylation analysis (n=237) by night shift work status. An asterisk (*) signifies a significant difference (p-value <0.05 using either χ^2^ test, Fisher’s exact test or 2-sided t-test) between non-night shift and night shift workers.

### Epigenome-wide methylation associations

DNA methylation at 298 CpG sites was found to be significantly different in night shift workers after FDR correction at the BH q<0.10 (Tables S2 and S3), 57 CpG sites significant at the BH q<0.05 (Table 2), and 10 CpG sites at the Bonferroni-corrected p<0.05 (Table 2).

**Table 2.**
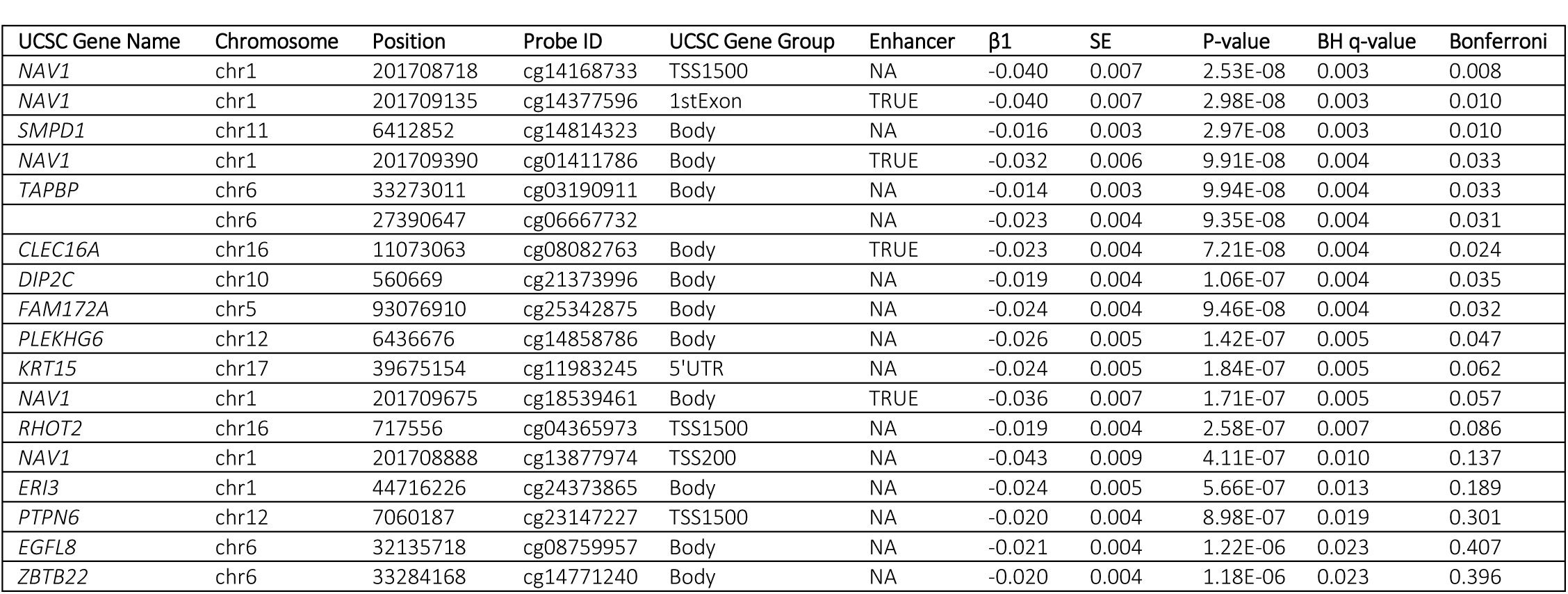
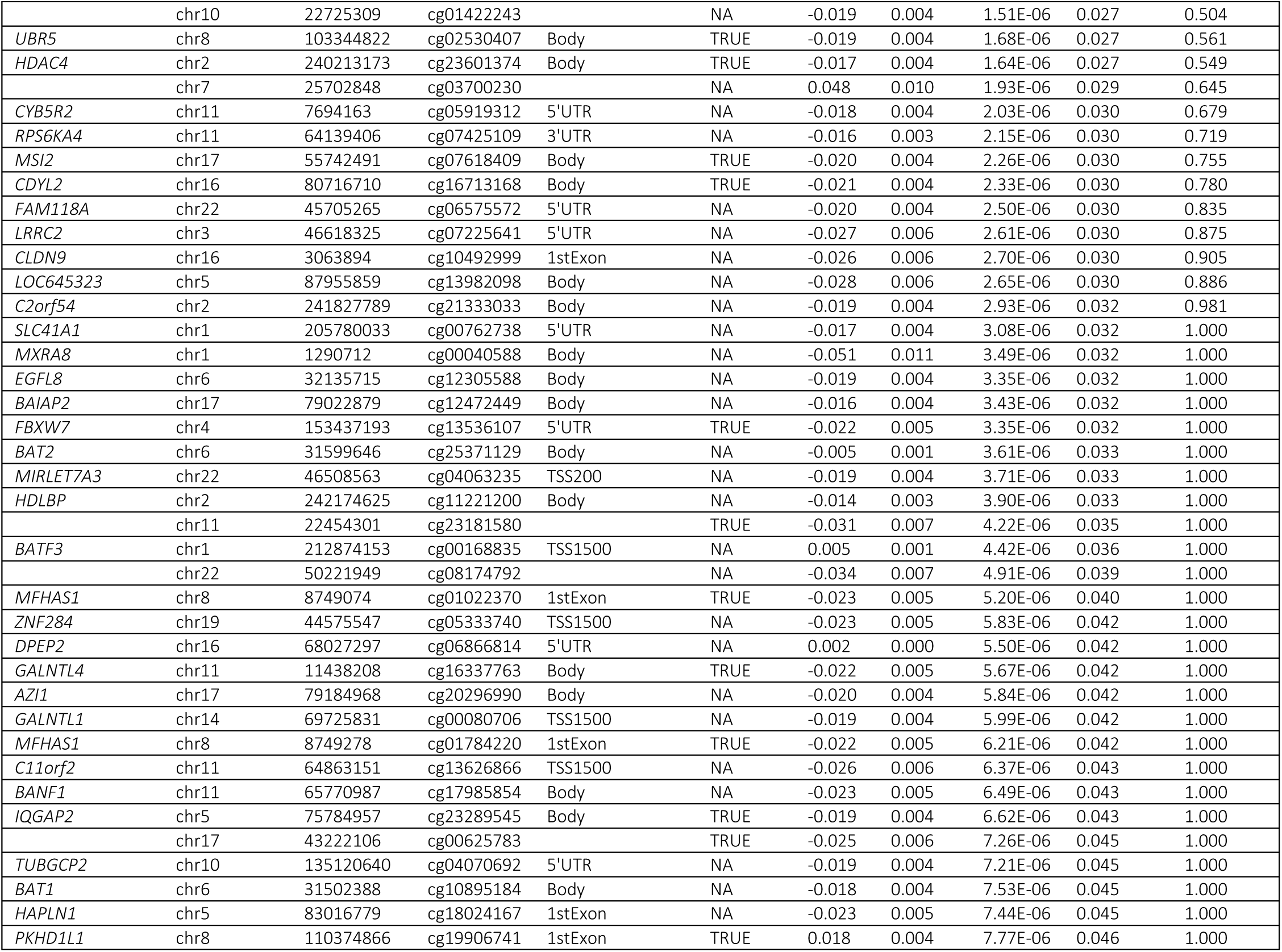
List of differentially methylated CpG sites in night shift workers compared to non-night shift workers after epigenome-wide analysis (BH q<0.05).

CpG sites for the *NAV1, SMPD1, TAPBP, CLEC16A, DIP2C, FAM172A,* and *PLEKHG6* genes had genome-wide significance after Bonferroni correction (p<0.05). The *ADAMTS10, CLEC16A, CTBP1, EGFL8, GNAS, HDAC4, HEATR2, KCNA4, KDELC2, MFHAS1, MXRA8, NAV1, PLXND1, UBR5, WNT5A,* and *ZBTB22* genes had multiple CpG sites represented in the results. There was an overall trend towards hypomethylation (Figure 1a). CpG sites for *NAV1, MXRA8, GABRG1, PRDM16, WNT5A,* and *FOXG1* were among the 10 sites with the most hypomethylation; CpG sites for *TDO2, ADAMTSL3, DLX2,* and *SERPINA1* were among the 10 sites with the most hypermethylation (Table S3).

**Figure 1a.**
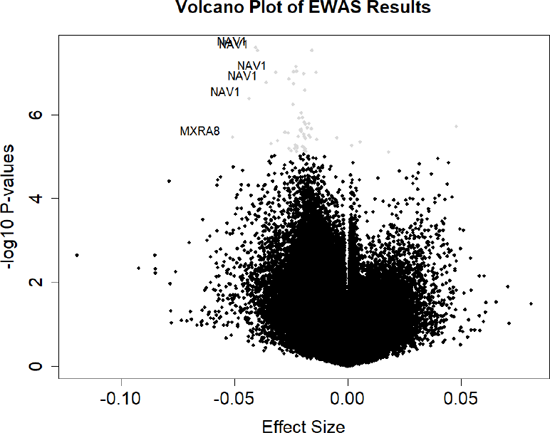
Volcano plot of effect size (beta coefficients, β1) and -log10 P-values of night shift EWAS, adjusted for maternal age, pre-pregnancy smoking, adversity score, sex of the infant, and estimated cell mixture. Gray dots signify CpG sites with BH q<0.05 and CpG sites with both absolute beta coefficients of 0.03 or greater and BH q<0.05 are labelled with UCSC gene names.

The Manhattan plot of the results indicated a number of differentially methylated sites that were distributed across the genome with some occurring in the same regions (Figure 1b). To more rigorously examine this finding, we employed a ′Bumphunter′ analysis and identified 6584 ′bumps′, with areas of the *NAV1, PURA, C6orf47,* and *GNAS* genes as DMRs (BH q<0.10)(Table 3). Of these, CpGs for the *NAV1* and *GNAS* genes were also differentially methylated in the CpG by CpG analysis (Table S3).

**Figure 1b.**
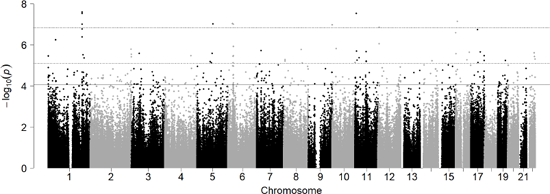
Manhattan plot of placental DNA methylation and night shift work EWAS, adjusted for maternal age, pre-pregnancy smoking, adversity score, sex of the infant, and estimated cell mixture. The dashed upper boundary line denotes p-value of 1.49×10-7 as the significance threshold after Bonferroni adjustment (p<0.05), the dashed middle boundary line denotes the p-value of 7.7×10-6 as the approximate significance threshold of BH q<0.05, and the solid boundary line at denotes the p-value of 8.8×10-5 as the approximate significance threshold of BH q<0.10.

**Table 3.**
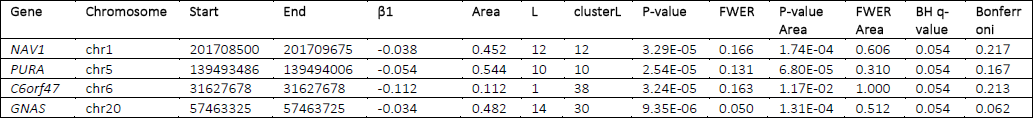
′Bumphunter′ results of significant DMRs (BH q<0.10).

A sensitivity analysis with GDM as a covariate shared many of the top CpG sites with the final results, suggesting GDM is not a main contributor to the findings (Table S4). An additional analysis evaluating GDM as the primary exposure shared no top genes with the EWAS results (BH q<0.10, data not shown). In another sensitivity analysis comparing the beta coefficients of those with *in utero* night shift work exposure only (n=37) to the beta coefficients of all night shift workers (n=53), the differences were small; only 1 CpG site, cg24373865, had an absolute difference in beta coefficients greater than 0.01, at 0.011. We also re-examined our results removing those with missing data on shift work and the findings were substantially similar (Table S5).

### Functional analyses

Comparing the 298 significant CpG sites (BH q<0.10) to the remaining 334,394 CpG sites, there was a higher frequency of top CpG sites within enhancer regions (χ^2^ = 13.48, df = 1, p-value = 0.0002). Because transcription factors (TFs) can bind to enhancer regions to alter gene expression, we assessed whether CpG methylation was associated with expression levels in nearby genes. The eQTM analysis found the expression of 18 genes to be associated with 14 CpG sites (p<0.05). Of these, the expression levels of *ACBD4* were associated with methylation in cg00625783 (β1=2.515, p-value=1.94E-05) and the expression levels of *KRT15* were associated with methylation in cg11983245 (β1=7.895, p-value=8.04E-05)(Table S6). For both of these genes, increasing methylation of the CpG sites was associated with increased gene expression. cg00625783 is not annotated to a gene but is located within an enhancer region and cg11983245 is annotated to the 5′ untranslated region (5′UTR) and 1^st^ exon of the *KRT15* gene. Methylation of cg11983245 was also associated (p<0.05) with increased *KRT19* (β1=4.404, p-value=3.87E-03) and *LINC00974* (β1=6.011, p-value=3.40E-02) expression levels.

We analyzed the top 298 CpG sites (BH q<0.10) for enrichment of KEGG pathways and GO-terms. The GO-terms "cell-cell adhesion”, "cell-cell adhesion via plasma-membrane adhesion molecules”, and "hemophilic cell adhesion via plasma membrane adhesion molecules” were found to be significant after FDR correction (BH<0.05)(Table S7). The top KEGG pathway results were "valine, leucine and isoleucine biosynthesis”, "mucin type O-glycan biosynthesis” and "melanogenesis”, but they were not significant after correcting for FDR (Table S7). Surprisingly, *PER1* was the only core circadian gene represented among the 298 CpG sites. However, we evaluated whether the 45 genes of the top 57 CpG sites exhibited circadian rhythmicity with the CircaDB expression database ^27^ and found 27 out of the 45 genes (60%) displayed rhythmic expression ^28^(Table S8). Of these genes, *BAIAP2, GALNTL1, HDLBP, NAV1,* and *TAPBP* displayed rhythmicity in mouse SCN tissue. To explore the physiological role of the Bonferroni (p<0.05) and ′Bumphunter′ significant genes, we queried the NHGRI-EBI GWAS catalog ^29^ for gene GWAS results with a p-value of 5×10^−8^ or less. The significant genes from the ′Bumphunter′ analysis were associated with traits such as BMI, blood pressure, the immune system, and autism spectrum disorder or schizophrenia in the GWAS catalog (Table 4).

**Table 4.** Query results of EWAS (Bonferroni p<0.05) and ′Bumphunter′ significant genes in NHGRI-EBI GWAS catalog ^29^. The gene name is the gene reported by the author(s) and/or the mapped gene and the listed traits are the traits reported by the author. GWAS results were filtered for significance with a p-value of 5×10^−8^.

| Gene | Number of Associated GWAS Studies | Reported Trait(s)<sup>29</sup> |
| --- | --- | --- |
| *NAV1* | 4 | BMI, BMI in physically active people, BMI adjusted for smoking, BMI in non-smokers, waist circumference |
| *SMPD1* | None |  |
| *TAPBP* | 1 | Autism spectrum disorder or schizophrenia |
| *CLEC16A* | 3 | Selective IgA deficiency, sum eosinophil basophil counts, eosinophil counts, eosinophil percentage of white cells, allergic disease (asthma, hay fever, or eczema) |
| *DIP2C* | 2 | Blood metabolite levels, uric acid levels |
| *FAM172A* | None |  |
| *PLEKHG6* | 2 | Colorectal cancer, primary biliary cholangitis |
| PURA | None |  |
| C6orf47 | 4 | Ulcerative colitis, inflammatory bowel disease, blood protein levels, autism spectrum disorder or schizophrenia, tuberculosis |
| GNAS | 7 | Platelet distribution width, diastolic blood pressure, systolic blood pressure, blood pressure, hypertension, renal function-related traits, BMI-adjusted waist circumference, DNA methylation |

## DISCUSSION

We identified a number of CpG sites exhibiting differential methylation associated with night shift work in newborn placental tissue. While the average absolute effect estimates for the 298 CpG site corresponded to a roughly 1.7% change in methylation, even a small change in methylation may have physiologically-relevant effects ^30^. The overall trend of hypomethylation with night shift work may be due to increased TF binding to DNA, leading to chromatin changes establishing the hypomethylated state^31^. Light at night (LAN) and night shift work can cause altered hormonal signaling and endocrine disruption; because hormone receptors can act as TFs, it is possible that circadian disruption causes increased hormonal signaling and increased TF binding.

Of the EWAS results, CpG sites for *NAV1* were consistently represented among the top results. The ′Bumphunter′ analysis also found a DMR in *NAV1*. In general, the functions of NAV1, particularly in the placenta, are not well characterized. *NAV1* is homologous to the *unc-53* gene in *C.elegans,* which plays a role in axonal migration ^32^. The mouse homolog also appears to play a role in neuronal migration; NAV1 is enriched in growth cones and associates with microtubule plus ends ^33^, and the deficit of *Nav1* causes loss of direction in leading processes ^34^. Research has also found increased embryonic lethality, decreased birthweight, and infertility in female offspring for Nav1^−/−^ mice ^35^, suggesting an important role for *Nav1* in fetal development and health. In eye tissue, *Nav1* was associated with mural cells, a precursor of pericytes, and may play a role in angiogenesis ^35^. In the embryonic retina, *Nav1* was downregulated in Math5^−/−^ mice, a TF affecting RGC differentiation, which suggests it may be associated with RGCs ^36^. Additionally, during embryonic development, *Nav1* was also found to be regulated by the TF PAX6, which has been implicated in sleep, brain and eye development, and metabolism, with Pax6^−/−^ mice having significantly lower *Nav1* mRNA expression in lens placode compared to wild type mice ^37^. When the CircaDB database of mouse tissue was queried, *Nav1* specifically displayed circadian rhythmicity in mouse SCN tissue (**Table S8**, JTK q<0.05). This suggests NAV1 may play a role in the mammalian SCN.

A DMR was also identified in *GNAS,* which is imprinted in the paraventricular nucleus of the hypothalamus and encodes the Gsa G-protein, which regulates cAMP generation and metabolism. *Gnas* is implicated in REM and NREM sleep and the browning of white adipose tissue for thermogenesis ^38^. Additionally, in a microarray analysis of retina samples from an *rd/rd* mouse model, *Gnas* was implicated in melanopsin signaling ^39^. Therefore, GNAS is likely important in integrating light and metabolic cues.

A possible limitation of this analysis is the moderate sample size of night shift workers (n=53). Additionally, the adjustment for cell-type heterogeneity is an estimation, so there is a possibility of residual confounding by cell type. Additionally, some of the women included as night shift workers did not have *in utero* exposure. Exposure to circadian disruption at different windows of development could have different magnitudes of effect. However, a sensitivity analysis of *in utero* night shift work exposure did not find large effect differences. Prior research has found that shift workers continue to have chronic health effects even after they switch to a day shift schedule. For example, researchers found that a history of shift work was associated with a decrease in cognitive ability that took 5 years or more after cessation of shift work to recover ^40^. This suggests recovery from regular shift work may take an extended period of time and a history of shift work may have a prolonged influence on health.

This is the first study to examine the epigenetic impacts of night shift exposure on placental methylation in humans. Methylation of placental tissue, an indicator of the *in utero* epigenetic landscape, reflects functional effects on the placenta, which can impact various aspects of fetal development, including neurodevelopment. The findings that the methylation of *NAV1* differed by night shift work exposure and that *Nav1* is rhythmically expressed in mouse SCN suggests NAV1 may play a developmental role in the human circadian system. Because the circadian system coordinates an array of physiological systems, alterations to circadian system development

Could affect immune response, sleep patterns, behavior, metabolism, and future health status. We have found night shift work to be associated with changes in methylation of placental tissue, which has implications for fetal development and future health. However, these findings may also be relevant for people who experience circadian disruption due to common exposures such as LAN.

## CONCLUSION

Night shift work is associated with differential methylation patterns in placental tissue. NAV1 may be an important component in the development of the human circadian system. Night shift work is a complex exposure encompassing altered hormonal signaling, eating and activity patterns, light exposure, and sleep patterns. Therefore, it is difficult to tease apart which aspects of night shift work contribute to which result. However, night shift work is a prevalent exposure in the workforce and, more generally, circadian disruption is a common facet of modern life. These findings warrant further investigation to evaluate the effects of *in utero* circadian disruption and the epigenetic programming of the circadian system.

## ACKNOWLEDGEMENTS

### Contributors

DACT designed and conducted the analysis. CJM led the cohort and collected the data used in the analysis. TME, AAB, MAD, KH, and JC provided code for the analysis and were involved in collecting the data. CJM, TME, AAB, and KEH helped with the conceptualization of the analysis and interpretation of the findings. DACT wrote the manuscript with the input and edits from all of the authors. All authors have reviewed and approved the final manuscript.

### Funding

This work was supported by the National Institutes of Health [NIH-NIEHS R01ES022223, NIH-NIEHS R24ES025145, NIH-NIEHS P01 ES022832, NIH-NIEHS T32 ES012870] and by the United States Environmental Protection Agency [US EPA grant RD83544201]. Its contents are solely the responsibility of the grantee and do not necessarily represent the official views of the US EPA. Further, the US EPA does not endorse the purchase of any commercial products or services mentioned in the manuscript.

### Competing interests

The authors have no competing financial interests to declare.

### Data sharing

The array data for RICHS can be found in the NCBI Gene Expression Omnibus (GEO) database with the number GSE75248 and the raw data for gene expression is available under the NCBI sequence read archive with the number SRP095910.

### Patient consent

Written informed consent was obtained from each participant for all aspects of the study.

### Ethics approval

RICHS and this analysis were approved by the Institutional Review Boards for the Women and Infants Hospital of Rhode Island and Emory University.

